# Radiotherapy Reshapes the 3D Genome in Medulloblastoma Cells

**DOI:** 10.64898/2026.10.09.757887

**Authors:** Dayna R. Challis, Richard Wilson, Sophie M. Navickas, Shannon N. Huskins, Iman Azimi, Adele F. Holloway, Kate H. Brettingham-Moore, Phillippa C. Taberlay

**Affiliations:** Menzies Institute for Medical Research, University of Tasmania, Hobart, Tasmania, Australia; South Australian Immunogenomics Cancer Institute, Adelaide University, Adelaide, South Australia, Australia; Central Science Laboratory, University of Tasmania, Hobart, Tasmania, Australia; Monash Biomedicine Discovery Institute, Monash University, Melbourne, Victoria, Australia; Tasmanian School of Medicine, University of Tasmania, Hobart, Tasmania, Australia

## Abstract

Radiotherapy is central to medulloblastoma treatment, yet the molecular and three-dimensional (3D) genome responses to irradiation remain poorly understood. Here, we integrated RNA sequencing, proteomics, and Micro-C to characterise the response of ONS-76 medulloblastoma cells to clinically relevant radiation exposures. A single 2 Gy dose induced widespread transcriptional and proteomic changes, including altered cell adhesion, migration, DNA replication, and ribosome biogenesis pathways. Integrated analysis identified LAMB3 and AKR1B10 as candidate mediators of radioresponse that remained elevated following fractionated irradiation. Micro-C profiling following single and repeated 2 Gy exposures revealed radiation-induced 3D genome remodelling. While global genome organisation remained largely stable, irradiation promoted shifts towards active A compartments, new TAD boundary formation, and transient boundary strengthening. Together, these findings provide the first genome-wide characterisation of radiation-induced 3D chromatin reorganisation in a medulloblastoma model and identify molecular and structural features associated with radioresponse.

## INTRODUCTION

Medulloblastoma is one of the most common malignant brain tumours in children and is typically treated using a combination of surgery, radiotherapy, and adjuvant chemotherapy (1–3). Following surgical resection, children over the age of three generally receive craniospinal irradiation, delivered in daily fractions of 1.6-2 Gray (Gy) over several weeks (2,3). Despite this intensive treatment, approximately 30% of patients experience disease recurrence, which is almost universally fatal (3,4). Radioresistance is thought to contribute to treatment failure and subsequent relapse, yet the mechanisms underlying this response remain poorly understood, with relatively few studies examining the molecular changes that occur in medulloblastoma cells following irradiation.

Cells respond rapidly to irradiation by altering gene transcription and protein abundance (5,6). High-throughput technologies such as RNA sequencing (RNA-seq) and mass spectrometry can capture these molecular changes at defined time points, providing a powerful strategy to uncover genes and pathways involved in radioresponse. DNA damage repair is a key contributor to radioresponse across cancers, alongside factors including the tumour microenvironment, heterogeneity, metabolism, and cell cycle dynamics (5,7–9). However, translating these mechanistic insights into effective radiosensitising strategies has proven challenging. Cetuximab, an EGFR-targeting monoclonal antibody approved by the FDA and TGA for the treatment of colorectal cancer and head and neck squamous cell carcinoma, remains the only molecularly targeted radiosensitiser approved for clinical use (9,10). In the context of brain cancer, most research in this field has focused on adult glioma models (11–14). Notably, Panizza *et al.* showed in glioma that proteomic, but not transcriptional, changes were the most accurate predictors of radiotherapy resistance (14), underscoring the value of multi-layered approaches for resolving treatment-induced cellular states. However, the molecular mechanisms underlying radioresponse in medulloblastoma are likely distinct from those of adult gliomas. Applying integrated proteogenomic approaches in medulloblastoma may therefore uncover disease-specific mechanisms of radioresponse.

Recent studies have also identified radiation-induced changes in 3D chromatin organisation in fibroblasts, lymphoblasts, and hepatocellular carcinoma cells; however, whether similar alterations occur in brain cancer cells remains unknown (15–17). Chromatin is organised hierarchically into large-scale chromosome territories (18), A/B compartments (19), topologically associating domains (TADs) (20), and fine-scale enhancer-promoter loops (21). Most previous studies have focused on the effects of radiation-induced DNA damage at the linear DNA level or have profiled local chromatin changes surrounding specific induced break sites (22,23). Importantly, only two studies have examined global changes in 3D chromatin organisation following radiotherapy (15,16). Sanders *et al.* assessed fibroblasts and lymphoblasts 30 minutes, 24 hours, and 5 days after a single 5 Gy dose of radiation using Hi-C (15). Notably, TAD boundary strengthening was observed after treatment, which may represent a mechanism to preserve 3D genome architecture (15). Similarly, Zhang *et al.* profiled hepatocellular carcinoma cells 30 minutes after a single 4 Gy dose of radiation using Hi-C and RNA-seq and found that TAD boundary disruption had the greatest effect on gene transcription (16). While these studies demonstrated clear radiation-induced changes in 3D chromatin structure, they neither used clinically relevant radiotherapy doses nor assessed repeated exposures (15,16). How 3D genome organisation responds to clinically relevant and repeated radiation exposure in medulloblastoma therefore remains unknown.

Here, we used a radiotherapy-naïve SHH-subgroup medulloblastoma cell line (ONS-76) to model radioresponse. RNA-seq and mass spectrometry following a single clinically relevant 2 Gy dose of radiation identified LAMB3 and AKR1B10 as candidate mediators of radioresponse. We then used Micro-C to profile genome-wide 3D chromatin organisation following single and repeated 2 Gy radiation exposures, capturing both transient and more sustained topological changes. Radiotherapy induced widespread compartmental shifts, predominantly from compartment B to compartment A, consistent with widespread redistribution towards a more active chromatin state. Together, these analyses indicate coordinated transcriptional, proteomic, and 3D chromatin remodelling in medulloblastoma cells following irradiation. By applying clinically relevant radiation doses and incorporating repeated exposures, our study addresses a key gap in the field and provides the first genome-wide assessment of radiation-induced 3D chromatin reorganisation in a medulloblastoma model.

## RESULTS

### Radiotherapy induces widespread transcriptional and proteomic changes in medulloblastoma cells

Before examining radiation-induced molecular changes, the baseline radiosensitivity of the ONS-76 cell line was compared to the well-characterised Daoy cell line using clonogenic survival assays and γH2AX immunofluorescence (Supplementary Fig. S1A,B) (24). ONS-76 cells displayed significantly reduced survival relative to Daoy cells following both 2 Gy (46% vs 68%; *p*>0.05) and 4 Gy (11% vs 27%; *p*<0.01) irradiation, consistent with a more radiosensitive phenotype (Supplementary Fig. S1). This increased sensitivity was further supported by γH2AX immunofluorescence following radiomimetic treatment, with ONS-76 cells showing greater and more persistent DNA damage than Daoy cells (Supplementary Fig. S1B). Fractionated treatment (15 × 2 Gy) generated ONS-76-RT cells with increased survival following 2 Gy irradiation (*p*>0.01) (Supplementary Fig. S1A). These findings confirm that ONS-76 cells not only represent an appropriate model to investigate early molecular responses to radiotherapy but can also develop a more radioresistant phenotype following repeated exposure, making them suitable for studying both acute and longer-term radiation responses.

To characterise these early molecular changes, ONS-76 cells were exposed to a single 2 Gy dose of radiation, and global changes in gene expression and protein abundance were assessed 24 hours post-treatment using RNA-seq and mass spectrometry. Overall, 601 genes were significantly upregulated, and 372 genes were significantly downregulated in the irradiated cells (Fig. 1A). *S*omatostatin receptor 5 antisense RNA 1 (*SSTR5-AS1*) was the most significant upregulated gene (FDR = 4.75 × 10⁻⁴; Log2FC = 2.16), while left-right determination factor 1 *(LEFTY1*) exhibited the greatest fold change (Log2FC = 2.35) (Fig. 1A). Similarly, 578 proteins showed significant changes in abundance (361 increased and 217 decreased) in the irradiated cells (Fig. 1B). Laminin subunit beta-3 (LAMB3) was the most significant upregulated protein (FDR = 5.58 × 10⁻³; Log2FC = 0.93), while TATA element modulatory factor 1 (TMF1) displayed the greatest fold change (Log2FC = 1.98), followed by syndecan-1 (SDC1) (Log2FC = 1.75) (Fig. 1B).

**Figure 1.**
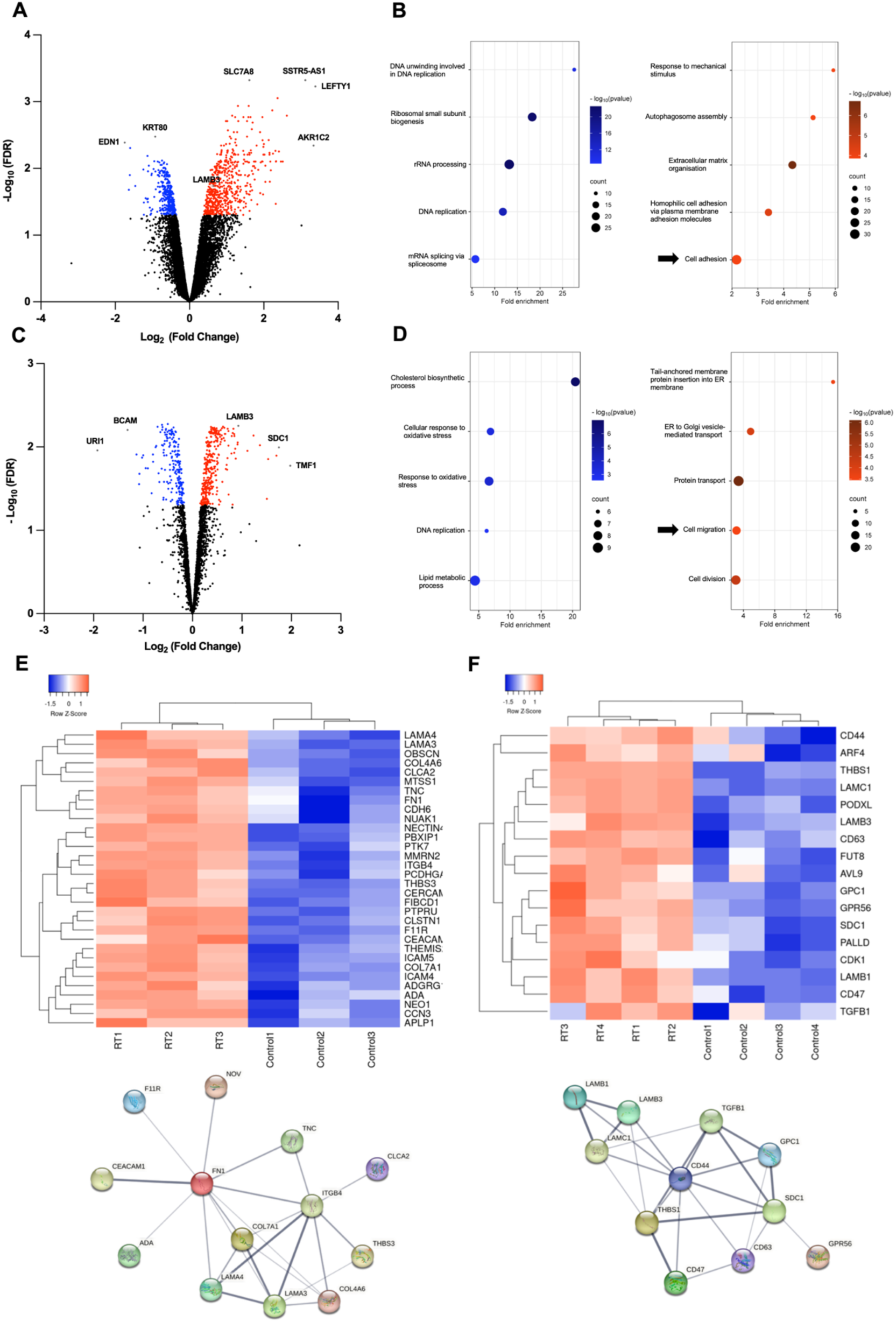
Radiotherapy induces transcriptional and proteomic changes associated with cell adhesion and migration. **A)** Volcano plot showing all differentially expressed genes following radiotherapy. ONS-76 cells were either left untreated or treated with a single 2 Gy dose of radiotherapy, and RNA-seq was performed 24 hours post-irradiation (n = 3). Significantly upregulated genes (FDR < 0.05) are highlighted in red, and significantly downregulated genes in blue. **B)** Top five Gene Ontology (GO) biological processes enriched among significantly upregulated (red) and downregulated (blue) genes. **C)** Volcano plot showing all proteins with altered abundance 24 hours after a single 2 Gy dose of radiotherapy, analysed by HPLC-MS (n = 4). Proteins with significantly increased abundance (FDR < 0.05) are shown in red, and those with decreased abundance in blue. **D)** Top 5 GO biological processes for proteins showing increased (red) or decreased (blue) abundance. **E)** Heatmap and STRING network of genes associated with the “cell adhesion” GO term. Interactions with a confidence score ≥ 0.4 are shown; disconnected nodes were excluded. Line thickness represents interaction confidence. **F)** Heatmap and STRING network of proteins associated with the “cell migration” GO term. Interactions with a confidence score ≥ 0.4 are shown; disconnected nodes were excluded.

To gain functional insight into these widespread transcriptional and proteomic changes, Gene Ontology (GO) enrichment analysis was performed to identify biological processes altered at both the transcript and protein levels following irradiation. Radiation rapidly activated cell adhesion, migration, and proteostasis programmes, while supressing DNA replication and ribosome biogenesis (Fig. 1C,D). Upregulated terms included cell adhesion and cell migration, alongside processes related to the unfolded protein response, including autophagosome assembly, ER-to-Golgi vesicle-mediated transport, and protein transport (Fig. 1C,D). In contrast, downregulated terms included DNA replication, rRNA processing, and oxidative stress pathways, reflecting suppression of biosynthetic and proliferative activity (Fig. 1C,D). Cell adhesion contained the greatest number of differentially expressed genes among the enriched GO terms (Fig. 1C), prompting us to further examine this response. A STRING (Search Tool for the Retrieval of Interacting Genes/Proteins) network of adhesion-associated genes highlighted Integrin Subunit Beta 4 (ITGB4), Fibronectin (FN1), and Collagen Type VII Alpha 1 Chain (COL7A1) as central nodes within the cell adhesion network (Fig. 1E). Concordantly, proteins associated with cell migration, including Syndecan-1 (SDC1), Laminin Subunit Beta-3 (LAMB3), and other laminins, were strongly upregulated following irradiation (Fig. 1D,F). Given these pathway-level changes across both datasets, we next integrated the transcriptomic and proteomic responses to identify individual molecules showing coordinated changes in gene expression and protein abundance.

### Multi-omics analysis identifies *LAMB3* and *AKR1B10* as candidate radiation response genes

To prioritise candidate molecules involved in the early radiation response, we integrated the RNA-seq and proteomics datasets. Forty-eight genes were significantly altered at both the transcript and protein levels (Fig. 2A; Supplementary Table S1), forming a cluster centred around RBM28 that was associated with ribosome biogenesis, as well as a second cluster associated with DNA replication and repair processes (Fig. 2A). Given that protein abundance may more closely reflect functional cellular changes (14), we next prioritised the 50 most significantly altered proteins, ranked by FDR (Fig. 2B), and intersected these with all differentially expressed genes (Fig. 2C). This identified three candidates altered across both datasets: *LAMB3*, *AKR1B10*, and *STMN1* (Fig. 2C). Of these, *LAMB3* and *AKR1B10* were consistently upregulated following irradiation (Fig. 2B, C) and remained elevated after fractionated radiotherapy (15 x 2 Gy) (Fig. 2D). *LAMB3*, encoding the β3 subunit of laminin-332, links directly to the cell adhesion and migration programmes identified above (Fig. 1), while *AKR1B10*, an aldo-keto reductase involved in aldehyde metabolism and oxidative stress regulation (25,26), may reflect the activation of broader stress-response pathways.

**Figure 2.**
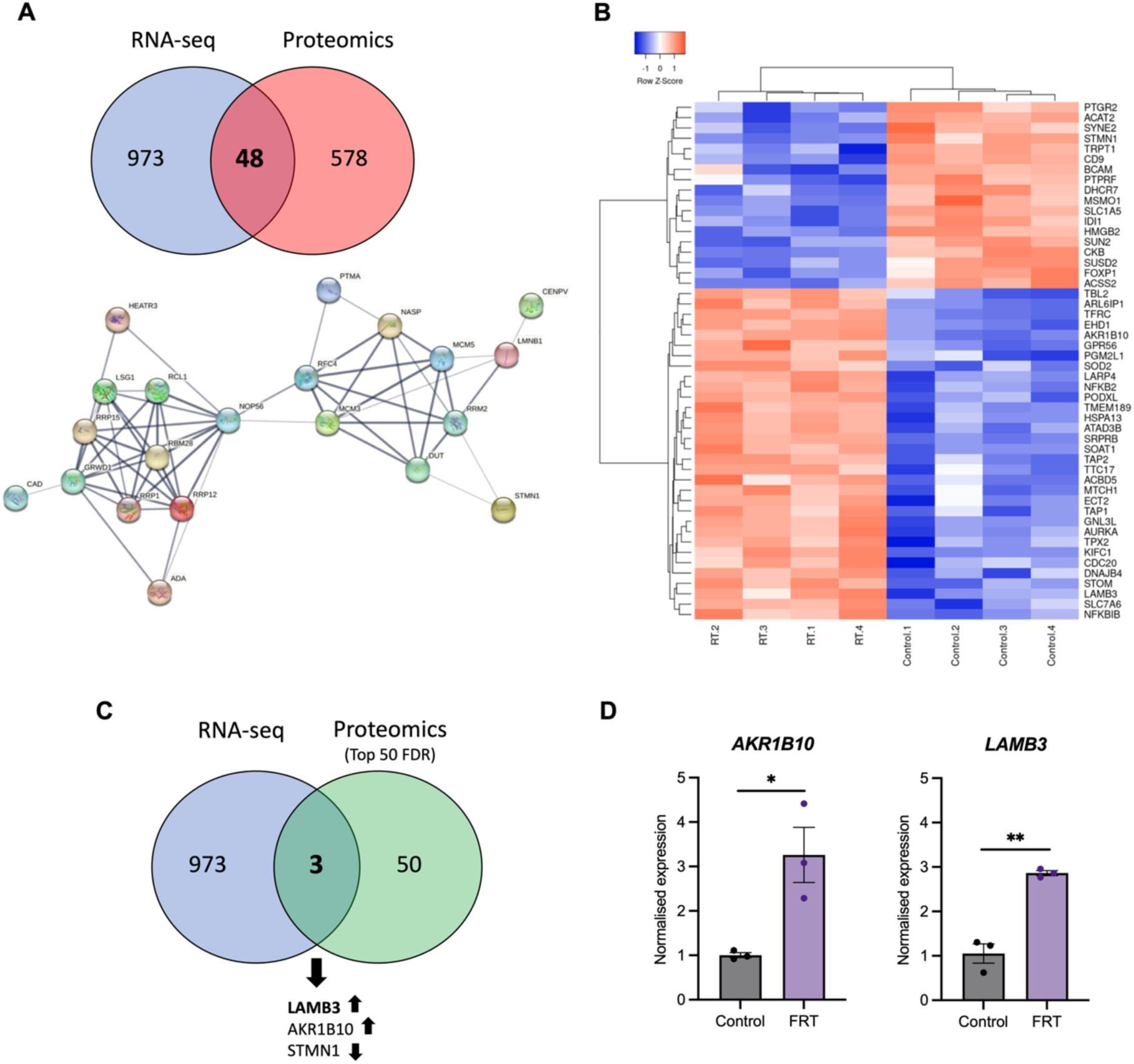
Transcriptome–proteome integration identifies key mediators of radiotherapy response. **A)** Venn diagram showing the overlap between all differentially expressed genes and proteins with significantly altered abundance following 2 Gy irradiation. STRING network visualising interactions between the shared genes; only interactions with a confidence score ≥ 0.4 are shown, and disconnected nodes were excluded. **B)** Heatmap of the top 50 proteins with altered abundance post-radiotherapy, ranked by adjusted p-value (FDR<0.0067). Red indicates increased abundance and blue indicates decreased abundance. **C)** Venn diagram intersecting the top 50 proteins with all differentially expressed genes, highlighting three shared candidates. **D)** Expression of *LAMB3* and *AKR1B10* following fractionated radiotherapy (15 × 2 Gy doses over 3 weeks). Expression levels were normalised to *B2M*, and fold change was calculated relative to the untreated control using the 2^-ΔΔCT method (mean ± SEM; n = 3). Significance was determined by Student’s t-test (*p<0.05, **p<0.01).

### Radiotherapy promotes a shift towards compartment A

A and B compartments reflect the spatial segregation of transcriptionally active euchromatin (A compartments) and inactive heterochromatin (B compartments) within the nucleus (27). Changes in compartment identity can influence gene expression (28) and may also affect susceptibility to DNA damage (29). To assess whether radiotherapy alters compartment organisation in ONS-76 cells, compartments were called at 1 Mb resolution, with the first eigenvector used to assign compartment identity (30). Most compartments remained stable following irradiation, with switches detected across approximately 10% of the genome 24 hours after a single 2 Gy dose and 8% of the genome five days after single or repeated doses (Fig. 3A). Notably, the majority of compartment switches were from compartment B to compartment A (B-A), indicating an overall shift towards active compartment status following irradiation (Fig. 3A).

**Figure 3.**
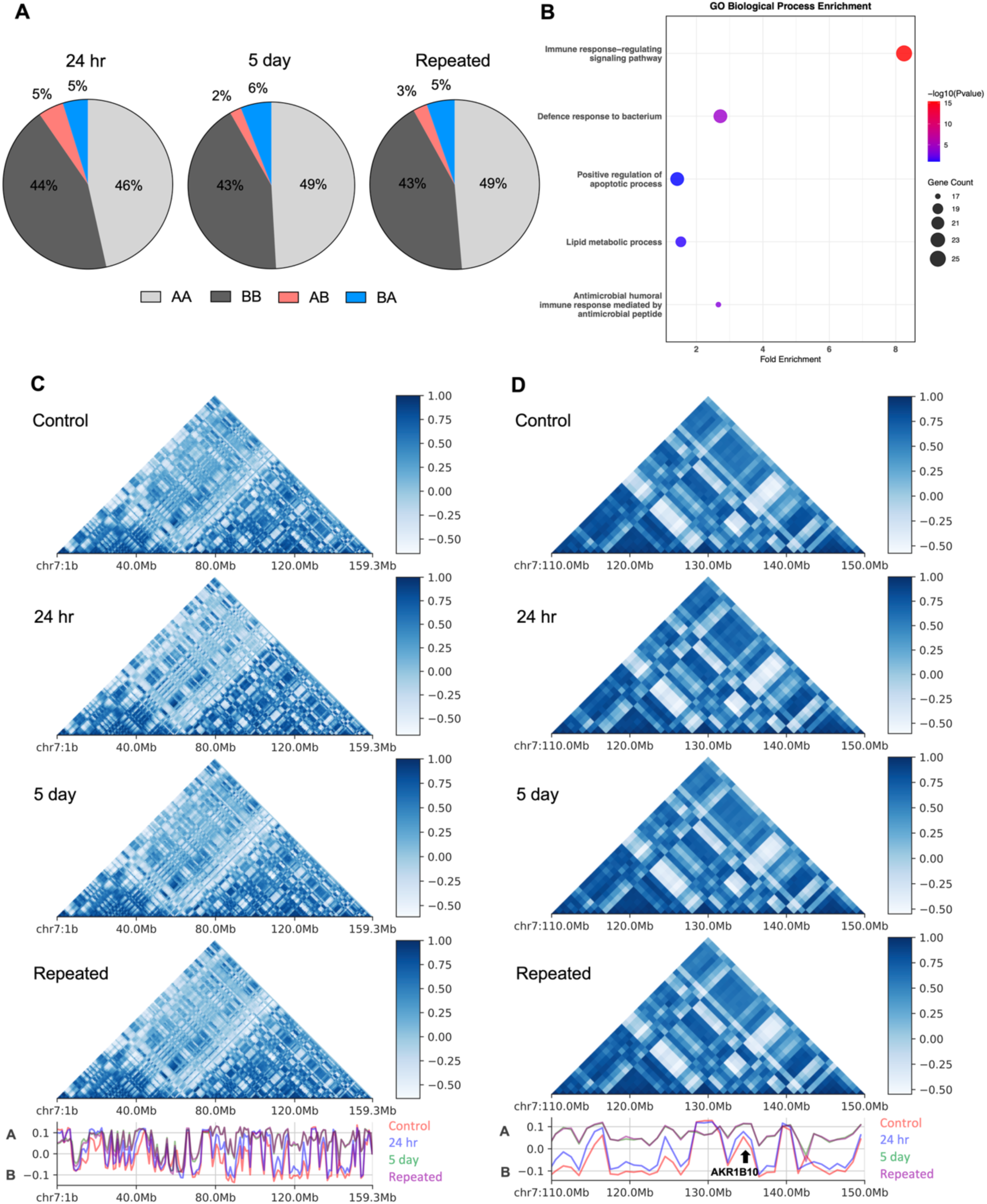
Radiotherapy induces progressive reorganisation towards A compartments. **A)** Overall changes in compartment identity in ONS-76 cells treated with a single 2 Gy dose radiotherapy (24 hr, 5 day) or repeated 2 Gy doses followed by 5 days of recovery (Repeated) were measured using Micro-C (n = 2). Compartments were called at 1 Mb resolution, and the proportions of stable (A-A, B-B) and switching (A-B, B-A) compartments were determined relative to the untreated control. **B)** Gene Ontology (GO) biological processes enriched among genes located within B-to-A (B–A) switch regions following repeated radiation doses. Genes were annotated to these regions and GO enrichment analysis was performed using the annotated gene sets. Dot size represents the number of genes in each GO term, and colour indicates the adjusted p-value (FDR). **C)** Contact maps showing compartment organisation across chromosome 7, binned at 1 Mb resolution. **D)** Magnified view of a 40Mb region on chromosome 7 binned at 1 Mb resolution. The first eigenvector from principal component analysis is plotted below each map; positive eigenvector values indicate A compartments, and negative values indicate B compartments.

To explore the potential functional relevance of these compartment shifts, we performed Gene Ontology (GO) enrichment analysis on genes located within regions that transitioned from compartment B to A following repeated radiation exposure (Fig. 3B). The most enriched biological processes included immune response, defence response to bacterium, and positive regulation of apoptotic process (Fig. 3B). These findings suggest that genomic regions gaining active compartment status after repeated irradiation are enriched for genes involved in immune-related and broader cellular stress responses.

Chromosome 7 exhibited some of the most pronounced compartment shifts following irradiation (Fig. 3C). A prominent B-to-A transition was observed near 112 Mb, where the control showed a negative eigenvector value, the 24-hour sample approached zero, and both the 5-day and repeated-dose samples showed positive eigenvector values (Fig. 3C), reflecting the broader B-to-A trend observed genome-wide (Fig. 3A). Given the identification of *AKR1B10* in our multi-omics analysis, we also examined a 20 Mb region of chromosome 7 (125–145 Mb) encompassing the *AKR1B10* locus (Fig. 3D). *AKR1B10* remained within compartment A across all conditions but showed increased positive eigenvector values after the 5-day and repeated-dose regimens (Fig. 3D). Similarly, *LAMB3* remained at the A/B compartment interface across all conditions (Supplementary Fig. S2).

### Irradiation induces transient TAD boundary strengthening and new boundary formation

Topologically associating domains (TADs) are self-interacting genomic regions that regulate gene expression by promoting enhancer–promoter communication within the same domain while limiting interactions across TAD boundaries (20). TADs are formed through loop extrusion, during which DNA is extruded through cohesin until blocked by boundary proteins such as CTCF (31). Despite extensive reports of TAD perturbation and the formation of new TAD-like structures in several cancers (32–34), their radiotherapy-induced dynamics remain largely undefined.

Here, the number and size distribution of TADs remained largely unchanged following single or repeated doses of radiotherapy (Fig. 4A,B). Across all conditions, 1841 TAD boundaries were shared, indicating that global TAD architecture is broadly stable. However, a distinct subset of boundaries was detected only after irradiation, suggesting locus-specific, radiation-induced remodelling (Fig. 4C). Notably, 64 boundaries undetected in untreated cells were present in every irradiated condition (Fig. 4D). A further 19 boundaries were detected at the 5-day time point following both single and repeated irradiation, while an additional 38 boundaries were unique to the repeated-dose condition (Fig. 4D).

**Figure 4.**
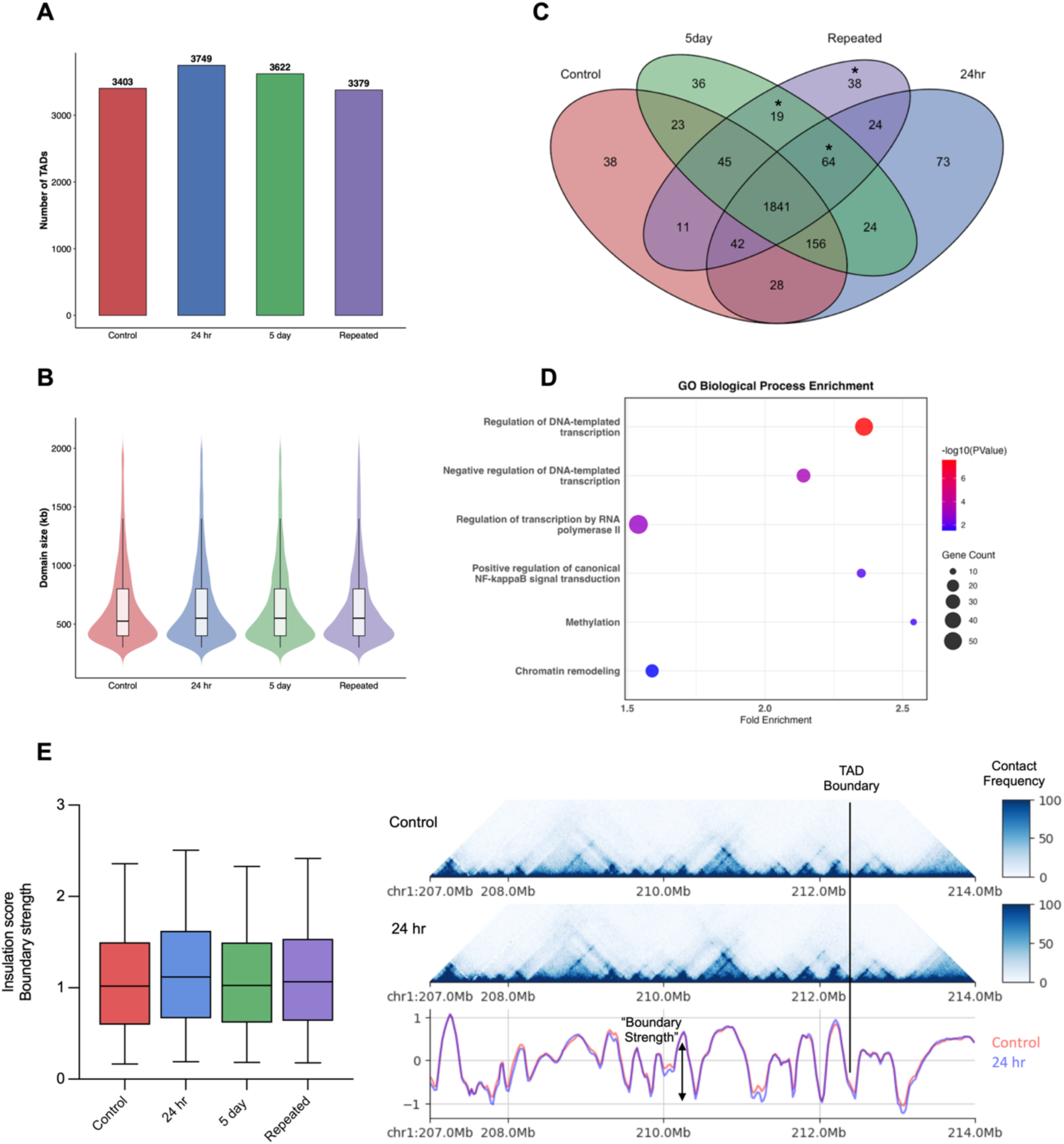
Irradiation induces formation of new TAD boundaries and transient strengthening of boundary insulation. **A)** Number of topologically associating domains (TADs) identified at 25 kb resolution in untreated ONS-76 cells (Control) and cells treated with a single 2 Gy dose of radiotherapy (24 hr, 5 day) or with repeated 2 Gy doses followed by 5 days recovery (Repeated), analysed using Micro-C. **B)** Distribution of TAD sizes across conditions. TAD size was calculated as the genomic distance between boundary pairs. **C)** Venn diagram showing the overlap of TAD boundaries between conditions. Asterisks denote the subset of boundaries used for downstream Gene Ontology (GO) enrichment analysis. **D)** GO biological processes enriched among genes located within gained boundary regions following irradiation. Dot size represents the number of genes in each GO term, and colour indicates the adjusted p-value (FDR). **E)** TAD boundary strength quantified using the insulation score method applied to 25 kb binned Micro-C matrices with a 500 kb insulation window. Boxplots show the median and upper and lower quartiles, with whiskers extending 1.5×IQR. Representative example of strengthened TAD boundaries 24 hours after irradiation. Contact maps depict a 7 Mb region of chromosome 1 binned at 25 kb resolution. Insulation scores (below) show a sharper dip at boundary positions in the 24 hr sample, indicative of increased boundary strength.

Genes located within these gained boundaries (marked by an asterisk) were annotated and GO enrichment analysis was performed (Fig. 4C,D). These genes were enriched for processes including regulation of DNA-templated transcription, regulation of transcription by RNA polymerase II, NF-κB signal transduction, methylation, and chromatin remodelling (Fig. 4D). Together, these results indicate that radiation-induced boundary formation preferentially occurs at loci containing genes involved in transcriptional control and chromatin regulation.

TAD boundary strength reflects the degree of insulation between neighbouring domains, with weaker boundaries permitting inter-TAD interactions and stronger boundaries providing greater insulation (35). Boundary strength was quantified using the insulation score method (36), applied to 25 kb binned matrices with a 500 kb window. Overall, TAD boundary strength showed a modest increase 24 hours after a single 2 Gy dose (median = 1.12) relative to untreated cells (median = 1.01), although this effect was not sustained at 5 days (Fig. 4E). In addition to these global changes, locus-specific boundary strengthening was evident. A representative 7 Mb region of chromosome 1 (207-214 Mb) illustrates this effect, where sharper decreases in insulation score indicate increased insulation following irradiation (Fig. 4E). These localised increases in insulation suggest that radiotherapy can selectively reinforce TAD boundaries, potentially constraining chromatin interactions at regulatory sites involved in the radiation response.

## DISCUSSION

Radiotherapy remains a key element of current medulloblastoma treatment; however, its use is limited by severe acute toxicities, long-term neurological complications, and the persistence of intrinsic or acquired radioresistance (4,37). Yet, despite its clinical importance, the mechanisms that shape the medulloblastoma radiation response are still poorly defined, and few studies have examined the global, multi-layered consequences of radiation in this disease. In this study, we interrogated the transcriptome, proteome, and 3D genome to characterise the early molecular and structural responses to clinically relevant radiation doses in ONS-76 medulloblastoma cells.

A substantial transcriptional and proteomic response was observed in ONS-76 cells after a single 2 Gy dose of radiation. Upregulated genes and proteins were enriched for biological processes associated with cell adhesion and migration, which may reflect early cytoskeletal remodelling rather than full epithelial-to-mesenchymal transition (EMT). The mesenchymal markers fibronectin (FN1) and N-cadherin (CDH2) were upregulated at the RNA and protein levels, respectively, while some epithelial markers, including SDC1 and several laminins, were also upregulated. This mixed pattern is consistent with partial EMT-like changes reported in other irradiated brain tumours (6,11), as well as functional evidence of increased migration after radiotherapy both *in vitro* and *in vivo* (38,39).

To move beyond pathway level patterns, integration of transcriptomic and proteomic datasets identified 48 genes altered at both the RNA and protein levels, with STRING network analysis revealing two major clusters associated with DNA replication and ribosome biogenesis. While changes in DNA replication are consistent with the response to radiation-induced DNA damage, ribosome biogenesis represents a less explored component of the radioresponse. The availability of pharmacological modulators of ribosome biogenesis provides an opportunity to investigate its potential role in radiosensitivity (40).

LAMB3 and AKR1B10 emerged as early radioresponse candidates, appearing among the top 50 most significantly altered proteins and showing upregulation at the RNA level. LAMB3 showed the most significant increase in protein abundance and encodes a subunit of laminin-332, a basement membrane protein involved in cell adhesion, migration, and survival (25,41), whereas AKR1B10 contributes to detoxification of cytotoxic carbonyls and lipogenesis (42,43). Both candidates have previously been linked to radioresistance in other cancers (26,44,45), and both genes showed further induction after three weeks of fractionated treatment in our study. This may reflect adaptation to repeated irradiation or selection of a subpopulation intrinsically expressing higher levels of these genes. Prior studies similarly report AKR1B10 upregulation in isogenic models of therapy resistance generated through repeated cytotoxic exposure (46,47). Together, these findings support functional investigation of both candidates in medulloblastoma, with selective AKR1B10 inhibitors providing a potential avenue for further study (48,49).

While radiation-induced mutations and chromosomal alterations have been extensively studied, the impact of radiotherapy on higher-order chromatin organisation remains poorly defined (15). Here, Micro-C was used to assess genome-wide changes in 3D chromatin structure and to characterise the chromatin context of candidate genes following radiotherapy. Both *LAMB3* and *AKR1B10* exhibited only minor compartment-level alterations, with *LAMB3* positioned at the A/B interface and *AKR1B10* located within compartment A. Approximately 8% of the genome underwent compartment shifts, and irradiated samples displayed a higher proportion of B-to-A transitions, consistent with a shift towards more transcriptionally active chromatin. However, only a small number of genes within these B-to-A regions displayed increased expression at 24 hours. Given that compartment identity influences but does not solely dictate transcriptional activity, and that later transcriptional time points were not assessed, these findings likely capture only a subset of compartment-associated regulatory changes. Genes located in regions gaining A-compartment identity after repeated radiation were enriched for immune response, defence response to bacterium, and regulation of apoptotic processes, suggesting that radiation-induced compartment changes occur at regions containing genes involved in immune and cellular stress responses. The reproducibility of these transitions across replicates further supports a coordinated rather than stochastic response to irradiation.

The number and size distribution of TADs remained largely unchanged following radiotherapy, consistent with findings in irradiated hepatocellular carcinoma cells (16). However, a small subset of boundaries was detected exclusively in irradiated samples, indicating locus-specific TAD remodelling. Genes located at these radiation-specific boundaries were enriched for NF-κB signalling, methylation, and chromatin remodelling pathways, suggesting potential regulatory effects. Integrating whole-genome bisulfite sequencing and ATAC-seq in future work would help clarify these mechanisms. TAD boundary strengthening was also observed after radiation exposure, consistent with other studies in UV- or X-ray-irradiated cells (15,17). Sanders et al. proposed that boundary reinforcement may result from increased recruitment of CTCF and cohesin to sites of DNA damage, thereby helping to preserve 3D genome integrity (15). In ONS-76 cells, boundary strengthening was evident 24 hours after a single dose of radiotherapy but not maintained at the 5-day time point, suggesting that this response is transient. Nevertheless, even transient strengthening may influence how cells respond to subsequent radiation exposures, which is particularly relevant given that clinical radiotherapy is delivered daily, at approximately 24-hour intervals (50).

Overall, this study provides a detailed characterisation of the radioresponse in ONS-76 medulloblastoma cells, demonstrating radiation-induced transcriptional, proteomic, and 3D structural changes. The identification of candidates such as LAMB3 and AKR1B10 highlights potentially important regulatory pathways, although further evaluation across additional medulloblastoma models and patient-derived systems will be essential to assess their broader relevance. While this multi-omics approach captures several key regulatory layers, complementary epigenetic and single-cell analyses will be necessary to more fully resolve chromatin dynamics and cellular heterogeneity following treatment. The therapeutic potential of candidate targets will also depend on effective delivery across the blood-brain barrier (BBB), although recent advances in materials science and nanotechnology provide potential avenues for brain-targeted drug delivery (51,52). Collectively, these findings show that both single and repeated radiation exposures can alter 3D chromatin architecture in medulloblastoma cells and identify genes, pathways, and topological changes that provide a foundation for future mechanistic and therapeutic studies.

## Acknowledgments

We thank Andrew McGrath, Samane Golmakani, and the team at the Holman Clinic, Royal Hobart Hospital, for their assistance with irradiating cell lines. This work was supported by Cancer Council Tasmania [B0027432], a donation from the Lions Club of Clarence, and the Australian National Health and Medical Research Council (NHMRC) [GNT1176494].

## Data Availability

All raw sequencing data generated in this study have been deposited to the NCBI Gene Expression Omnibus (GEO; https://www.ncbi.nlm.nih.gov/geo/) under accession numbers GSE349882 and GSE349548. Proteomics data is available via ProteomeXchange with identifier PXD085046. Additional data supporting the findings of this study are available from the corresponding author upon request.

## METHODS

### Cell culture and treatments

The ONS-76 (IFO50355; CellBank Australia) and Daoy (HTB-186; ATCC) medulloblastoma cell lines, both belonging to the sonic hedgehog (SHH) molecular subgroup, were used in this study. ONS-76 cells are TP53 wild-type and were derived from a 2-year-old female, whereas Daoy cells are TP53-mutant and were derived from a 4-year-old male. ONS-76 cells were maintained in RPMI 1640 supplemented with 2 mM L-glutamine, 10% fetal bovine serum (FBS), and 100 U/mL penicillin/100 μg/mL streptomycin (1% pen-strep). Daoy cells were cultured in MEM supplemented with 2 mM L-glutamine, 1 mM sodium pyruvate, 10% FBS, and 1% pen-strep. Cells were passaged at 80% confluency approximately twice weekly.

Cell irradiations were performed as described previously (5). To assess DNA damage by immunofluorescence, cells were treated with the radiomimetic phleomycin (1 μg/mL; Sapphire Biosciences) for 1 hour, a dose optimised to induce DNA damage comparable to 2 Gy irradiation (6,53,54). After treatment, cells were washed with PBS and incubated in fresh medium for the indicated recovery period.

### Clonogenic assay

The clonogenic assay was performed as previously described (5). Briefly, 1000 cells were plated into wells of a 6-well plate following 0, 2, 4, or 8 Gy irradiation. After 8 days of colony growth, medium was removed, and cells were washed in 1 ml of PBS. Colonies were fixed for 5 minutes in 800 μL of 3:1 methanol to glacial acetic acid. Fixative was removed, wells were air dried, and cells were stained with 1.0% methylene blue in 50% ethanol for 30 minutes. Percentage cell survival was calculated as the number of colonies post-treatment relative to the corresponding 0 Gy control.

### Immunofluorescence assay

γH2AX immunofluorescence was performed as previously described (6). Images were captured using a Nikon Eclipse Ti-2 Inverted Microscope at 40 x magnification with an exposure time of 100 ms. Deconvolution was applied to all images and Fiji Image J software (v 2.9.0) was used for image analysis (55). Images were converted to 8-bit grey scale and foci were quantified using the analyse particles function. Foci were quantified in at least 25 cells for each replicate (≥75 cells per condition).

### RNA-sequencing

RNA was extracted using TRI reagent®. RNA samples underwent clean-up using the RNA Clean & Concentrator™-5 kit (Zymo Research) following the manufacturer’s instructions. Library preparation and RNA-sequencing were performed at the Australian Genome Research Facility (AGRF). The Illumina NovaSeq 6000 platform was used to generate 100 bp single end reads.

Data was analysed using the online platform Galaxy (https://usegalaxy.org/). Firstly, the quality of raw Illumina sequence reads was assessed using FastQC (v 0.11.9) (56). To ensure data was suitable for downstream analysis, the first base was removed from each read across all samples using Trimmomatic (v 0.38.0) (57). HISAT2 (v 2.2.1) (58) was used to map reads to the human reference genome build hg38 and featureCounts (v 2.0.1) (59) was used to convert aligned reads into gene counts. Files were combined into a single count matrix and annotateMyIDs (v 3.12.0) was used to assign gene IDs. Finally, differential expression analysis was performed using edgeR (v 3.34.0) (60). Note, genes with less than 1 count per million (CPM) in at least four samples were filtered out. Volcano plots were created using GraphPad Prism (v 9.5.1) and heatmaps were generated using heatmapper (http://heatmapper.ca/). Upregulated and downregulated gene lists were also uploaded to DAVID for functional annotation (61).

### RT-qPCR

RNA was converted to cDNA using the the iScript™ cDNA Synthesis Kit (Bio-Rad). cDNA was amplified on the QuantStudio™ 3 real-time PCR system (Thermo Fisher Scientific) using Luna® Universal qPCR Master Mix (New England BioLabs) according to the manufacturer’s instructions. Primer sequences can be found in supplementary table 2. Cycling conditions were 95°C for 60 seconds, followed by 40 cycles of 95°C for 15 seconds and 60°C for 30 seconds. Melt analysis was used to confirm the presence of a single PCR product.

### Proteomics profiling

Protein was extracted from untreated and irradiated ONS-76 cells using denaturation buffer (7 M urea, 2 M thiourea, 40 mM Tris, pH 7.5) supplemented with protease inhibitors (cOmplete Mini EDTA-free cocktail, Merck). Protein concentration was determined by Bradford assay, and 30 µg aliquots were reduced, alkylated, and digested with 2 µg proteomics-grade trypsin/rLysC (Promega) using the SP3 method (62). Peptides were desalted with C18 ZipTips (Merck), dried by SpeedVac, and reconstituted in 12 µl HPLC loading buffer. Approximately 1 µg peptide was analysed by nanoflow HPLC–MS/MS on an Ultimate 3000 nano RSLC system coupled to a Q-Exactive HF mass spectrometer with a nanospray Flex ion source (Thermo Fisher Scientific) controlled by Xcalibur (v 4.3). Peptides were preconcentrated on a 20 mm × 75 µm PepMap 100 C18 trap column and separated on a 250 mm × 75 µm PepMap 100 C18 analytical column (300 nL/min, 45 °C) using a 120 min gradient. MS parameters were spray voltage 2.0 kV, S-lens RF level 60, and heated capillary 250 °C. MS1 spectra (390–1240 m/z) were acquired in profile mode at 120,000 resolution with an AGC target of 3 × 10^6, followed by data-independent MS2 acquisition across 26 × 25 amu windows (397.5–1027.5 m/z, 1 amu overlap). MS2 spectra were acquired in centroid mode at 30,000 resolution with an AGC target of 1 × 10^6, maximum IT 55 ms, and normalized collision energy 27.

DIA-MS raw files were processed using Spectronaut (v 15.0; Biognosys AB) as previously described (6). Spectronaut reports were imported into Perseus (v1.6.15.0) (63) for downstream statistical analysis. Missing values (<1% of proteins) were imputed using default settings. Differential protein abundance between untreated and irradiated samples was assessed using a Student’s t-test with FDR set to 5% and s₀ = 0.1 to filter out proteins with minimal fold changes. Data visualisation included volcano plots generated in GraphPad Prism (v 9.5.1) and hierarchical clustering of z-score-normalised data using Heatmapper (http://heatmapper.ca/). Protein lists were also uploaded to DAVID for functional annotation (61).

### Micro-C

Micro-C libraries were prepared using the Dovetail® Micro-C Kit according to the manufacturer’s instructions. Briefly, chromatin was fixed with disuccinimidyl glutarate (DSG) and formaldehyde in the nucleus. The cross-linked chromatin was then digested in situ with micrococcal nuclease (MNase). Following digestion, the cells were lysed with SDS to extract the chromatin fragments, and the chromatin fragments were bound to Chromatin Capture Beads. Next, the chromatin ends were repaired and ligated to a biotinylated bridge adapter followed by proximity ligation of adapter-containing ends. After proximity ligation, the crosslinks were reversed, the associated proteins were degraded, and the DNA was purified then converted into sequencing libraries using Illumina-compatible adaptors. Biotin-containing fragments were isolated using streptavidin beads prior to PCR amplification. Libraries were sequenced at the AGRF on the Illumina NovaSeq platform to generate a minimum of 300 million 2 x 150 bp read pairs per library. Micro-C was performed on two biological replicates per condition, generating a minimum of 360 million valid read pairs per condition which provided sufficient coverage for TAD analysis at 25 kb resolution.

Data was processed using the pipeline provided by Dovetail™ Genomics available on GitHub (https://micro-c.readthedocs.io/en/latest/index.html). Data quality was confirmed by assessing the fraction of cis-long range contacts for each library. To convert pairs files into contact matrices Juicer Tools (v 1.22.01) (64) was used and Knight-Ruiz (KR) normalisation was applied (65). Firstly, A/B compartments were called at 1 Mb resolution using FAN-C (v 0.9.24) (30). Eigenvectors (first principal-component) and compartment strength were determined using the appropriate FAN-C arguments. Next, TADs were called at 25 Kb resolution using Juicer arrowhead (64), and TAD boundaries were identified using the insulation score method (36). Insulation scores were calculated on 25 kb binned matrices using a 500 kb insulation square size.

## SUPPLEMENTAL FIGURES AND TABLES

**Supplementary Figure 1.**
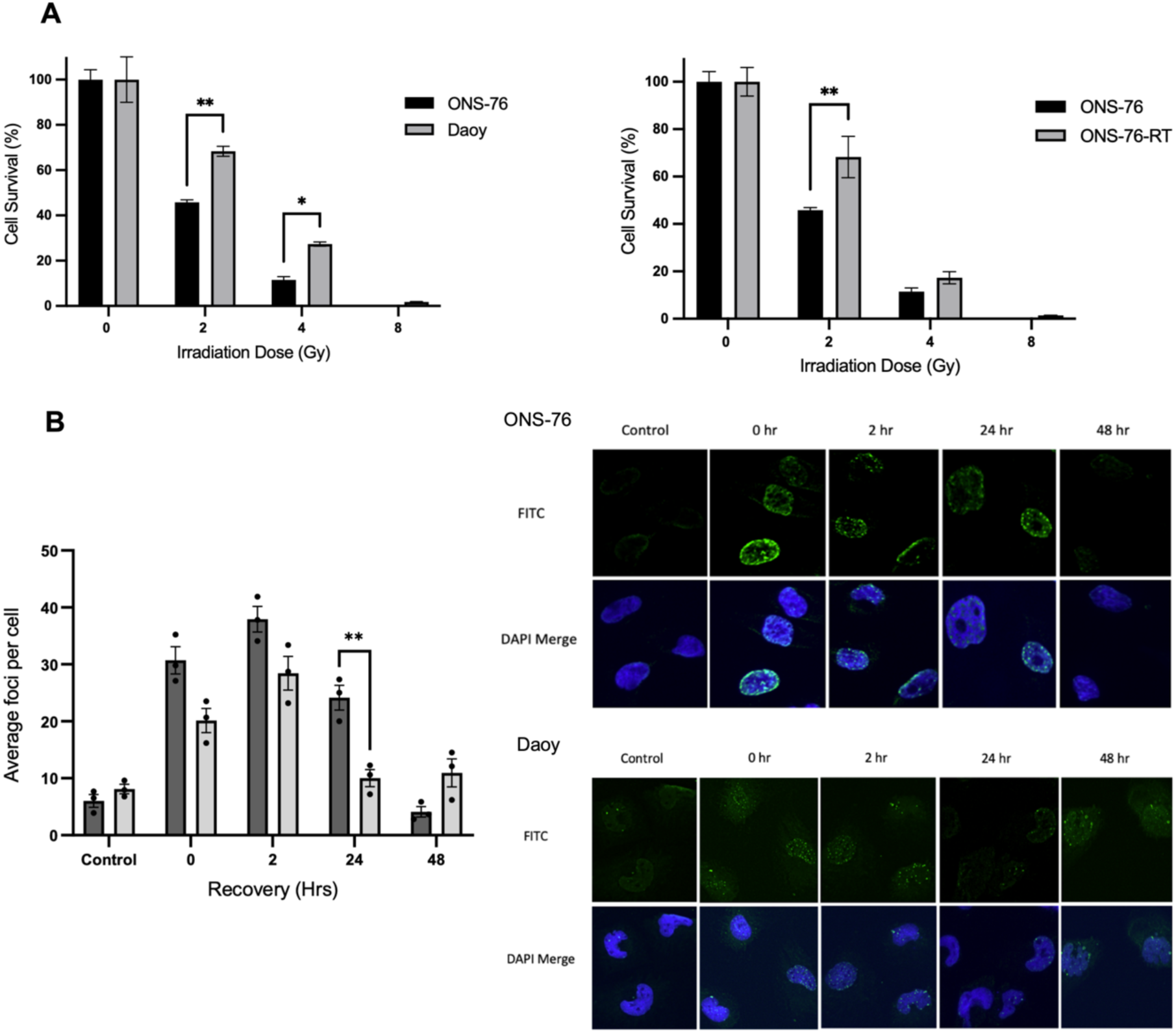
Radiosensitivity of ONS-76 and Daoy medulloblastoma cells. **A)** Clonogenic survival of ONS-76 and Daoy cells following the indicated doses of radiotherapy, and parental ONS-76 and ONS-76 cells exposed to three weeks of fractionated radiotherapy (ONS-76-RT; 15 × 2 Gy). Colonies were counted after seven days of growth and percentage cell survival was calculated relative to the untreated control (0 Gy) (mean ± SEM; n = 3). Significance was determined using two-way ANOVA with Sidak’s multiple comparisons test, *p>0.05, **p>0.01. **B)** γH2AX immunofluorescence in ONS-76 and Daoy cells following treatment with the radiomimetic phleomycin (1 μg/mL for 1 hour) and recovery for the indicated time periods. γH2AX foci were quantified in ≥75 cells per condition (mean ± SEM; n = 3). Representative images show γH2AX (FITC; green) and DAPI-stained nuclei (blue) at each recovery time point. Significance was determined using two-way ANOVA with Sidak’s multiple comparisons test, **p>0.01.

**Supplementary Figure 2.**
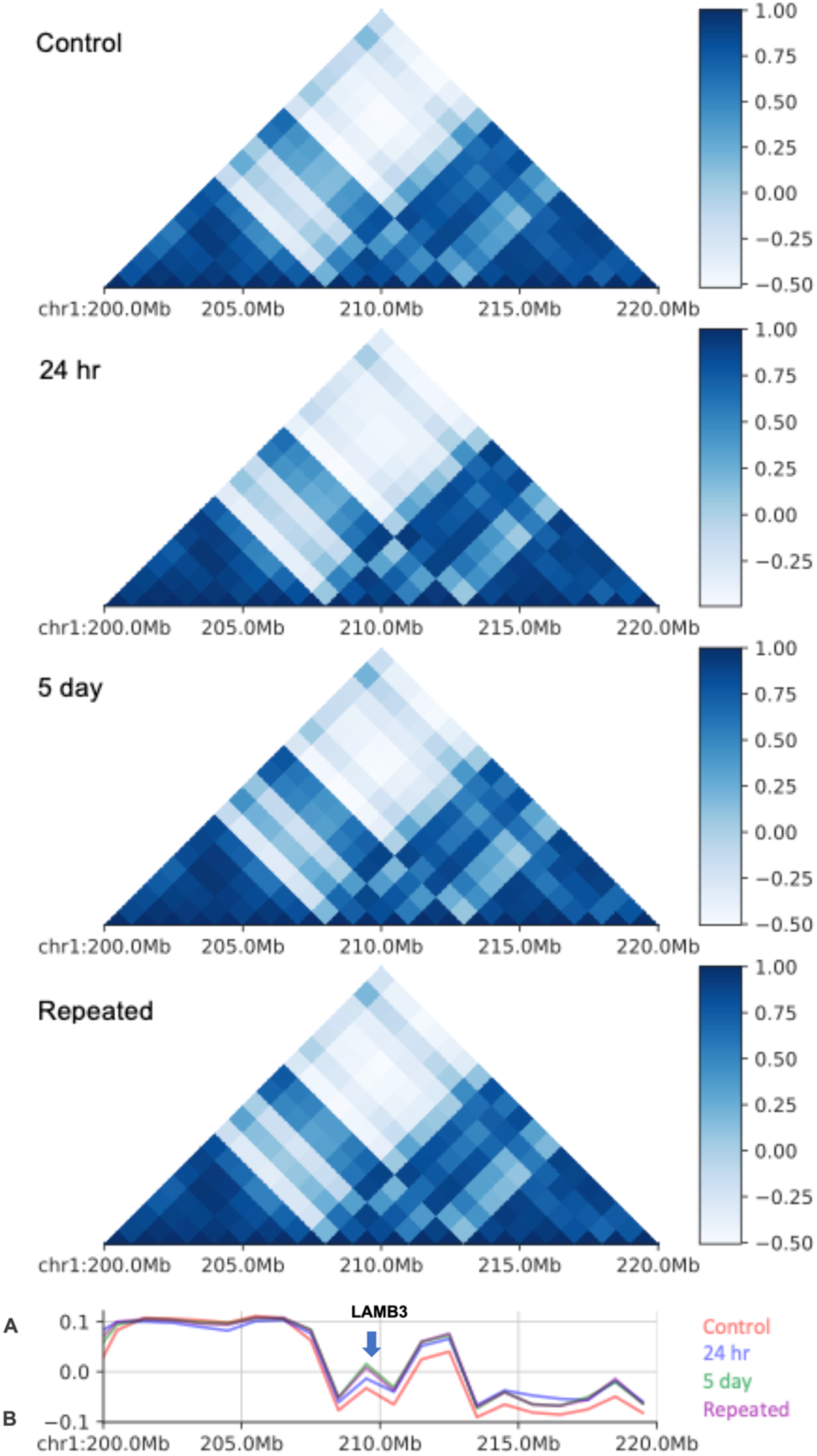
LAMB3 is located at the A/B compartment interface. Untreated ONS-76 cells (Control) and cells treated with a single 2 Gy dose of radiotherapy (24 hr, 5 day) or repeated 2 Gy doses of radiotherapy followed by 5 days recovery (Repeated) were analysed using Micro-C. Contact maps show a 20 Mb region of chromosome 1 binned at 1 Mb resolution. The first eigenvector from principal component analysis is plotted below. Positive eigenvector values indicate active A compartments, while negative values indicate inactive B compartments.

**Supplementary Table 1.**
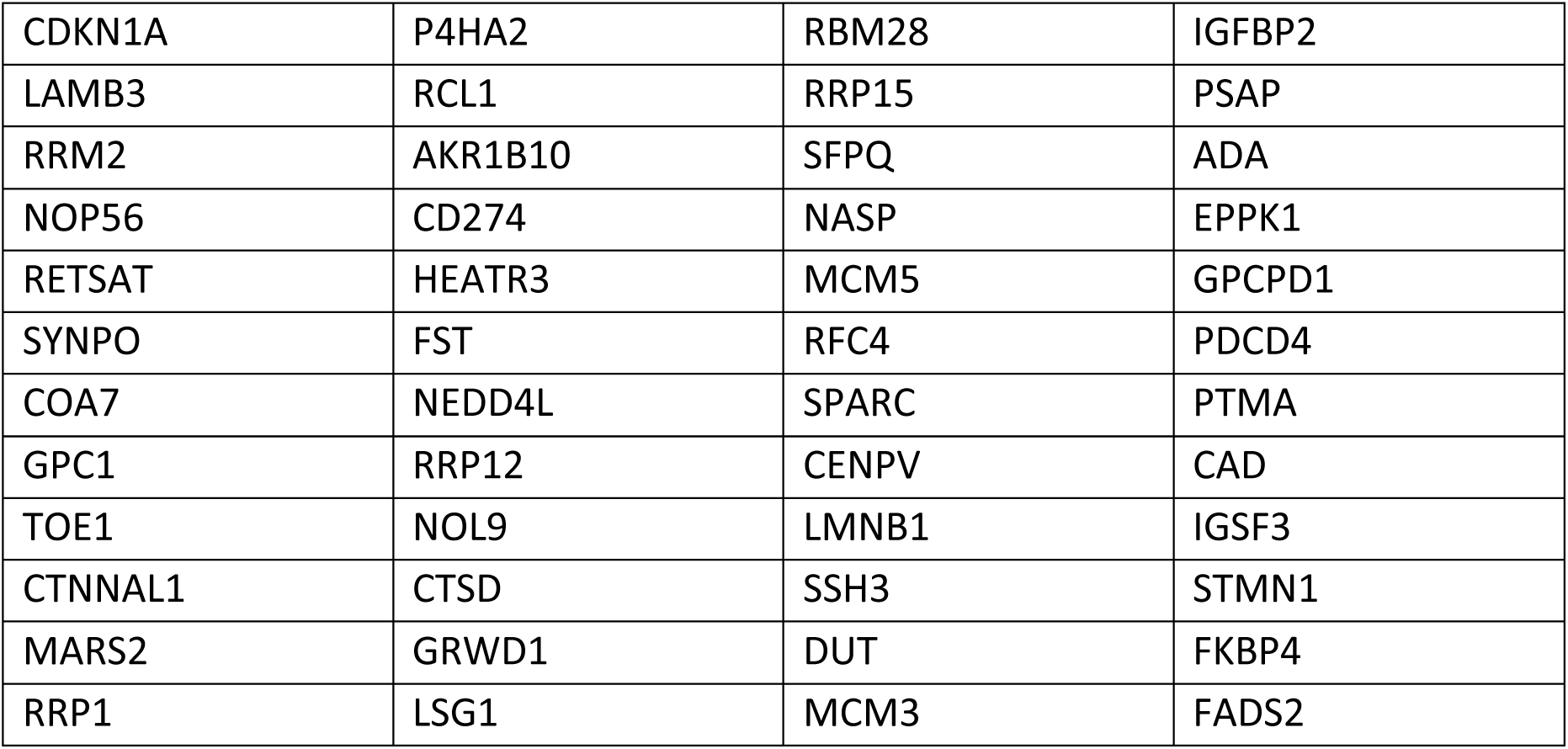
List of 48 genes shared between transcriptomics and proteomics datasets.

**Supplementary Table 2.**
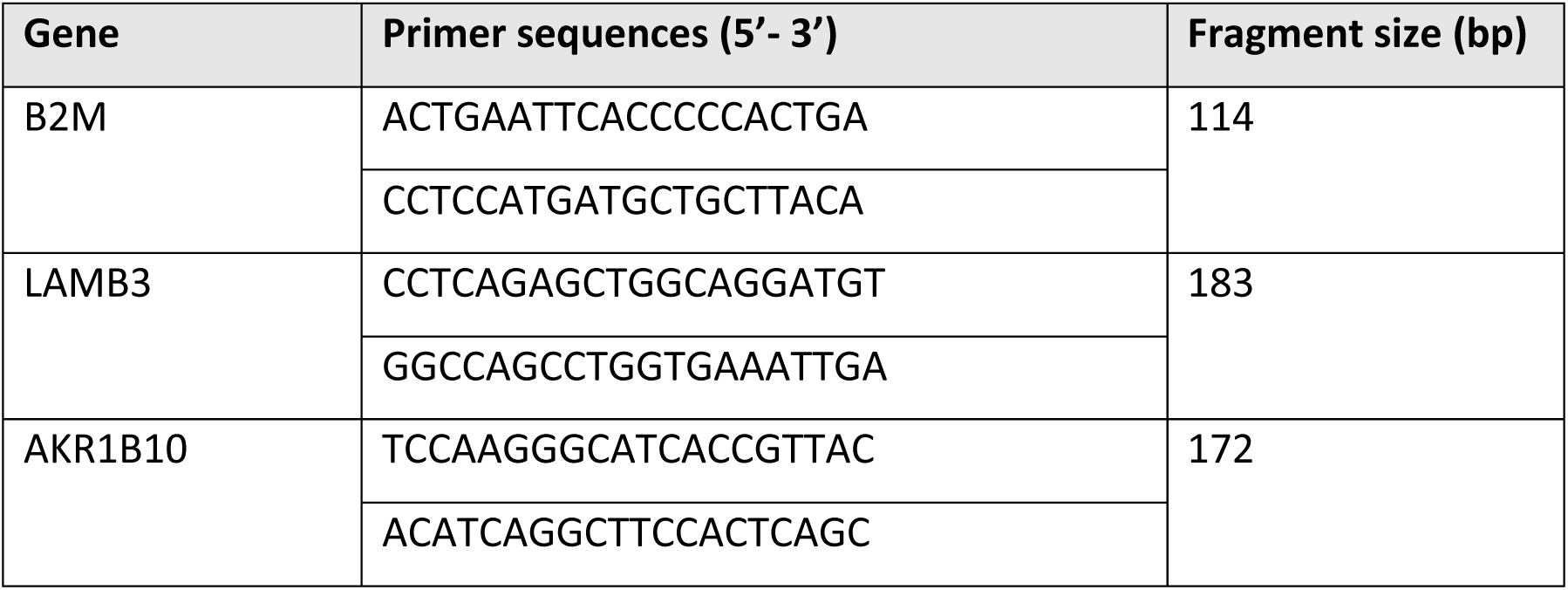
Primers for RT-qPCR analysis.

